# Physiological response to slalom water skiing: A case study of a sit-skier with paraplegia

**DOI:** 10.1101/858902

**Authors:** David Suárez-Iglesias, Carlos Ayán Pérez, José Antonio Rodríguez-Marroyo, José Gerardo Villa-Vicente

## Abstract

Recreational and competitive slalom waterskiing is popular among those with spinal cord injuries. People with paraplegia can practice on the slalom course using a sit-ski. A slalom run consists of a boat towing the sit-skier through a set of buoys and normally begins with a deep-water start. Despite its popularity, very little is known about the physiological aspects of the sit-skier's preparation. We examined the internal training load (TL) experienced by a sit-skier with paraplegia while learning and improving the slalom deep-water starts, executed with both the traditional technique and an alternative method. The TL was determined by means of heart rate (HR) and session rating of perceived exertion (sRPE) methods. The percentage of maximal heart rate values ranged from from 63.2% to 81.3% during deep-water starts. Training sessions were performed most of the time below the ventilatory threshold and tended to be qualitatively described as hard. A moderate but non-significant correlation existed between HR and sRPE-based methods. We also found a significant decrement in handgrip strength after practice. These findings indicate that the intensity of training experienced by our sit-skier was moderate in terms of physiological internal load during an adaptive slalom waterskiing training program.

## Introduction

In recent years, adaptive athletes have become more active and their participation exponentially has increased in minor and major events. Accordingly, there is increasing concern that a high level of physical conditioning and knowledge of training methodology are required when preparing athletes with motor disabilities for peaking [1]. However, a limited number of studies has examined adaptive sports performance training, the vast majority have been specifically focused on wheelchair sports [2,3]. In contrast, much uncertainty still exists about the Paralympic sporting disciplines outside the Paralympic Games. This is the case of adaptive waterskiing, a sport that offers recreational opportunities for individuals with spinal cord injuries (SCI) and varying degrees of abilities who are unable to stand [4].

The main event within sit-skiing is the discipline of slalom, in which developing a repeatable technique that creates the minimum load during deep-water starts (DWSs), and the ability to perform accurate DWSs, are crucial for the sit-skier’s success [5]. Deep-water starting can be characterized as a static exercise where the athlete resists high tow rope loads of ~2.0 – 2.5 times body weight, as reported in standing slalom waterskiers [6,7]. The conventional DWS maneuver not only presents complex technical aspects but may also be physically exhausting for individuals who are dependent on upper-body muscular strength and endurance due to the large isometric component. Thus, various DWS methods are taught, but whether a sit-skier is starting from the water independently, using an instructor-assisted start technique (e.g., arm-lock, straddle or side-skier methods), or using learning aids (e.g., “*deep V*” handle, “*Edge Triple Bar*”, “*boom*”) [8], there is always the need to reduce the required grip force and lessen the impact of fatigue [9]. Up to now, most studies in the field of standing waterskiing have focused on the biomechanical characteristics and physiological demands of slalom [5,7,10–13] and have adequately described a strength and conditioning program [14]. Despite its relevance from a performance point of view, the nature of the internal training load (TL) when starting and performing the repeated movements of slalom waterskiing remains unclear.

If coaches could have access to information regarding the TL that sit-skiers experience while learning and improving the slalom DWS and the several repeated movements for skiing to and around each of the 6 turn buoys, the training process of these skills could be improved, and specific training considering the characteristics of each athlete could be carried out. This goal could be achieved by registering data from parameters usually monitored in adaptive sports such as heart rate (HR) or rate of perceived exertion (RPE) [3]. Also, training zones are particularly useful in the case of individual sports to guide the training process [15,16]. These HR and RPE-based methods could provide a point of reference for coaches to compare the performance of their athletes with the data of participants in other studies, as well as to monitor the skiers themselves at different times of the season.

Finally, handgrip strength has been previously characterized in diverse adaptive athletes as a measure of physical performance [17–20], and it would be of interest to observe the workload intensity-related decline in handgrip strength during adaptive waterskiing practice. In this regard, athletes with SCI have been shown to commonly face overuse injuries to their upper limbs [21], and considering the stress levels imposed on skiers' upper bodies during the DWS [14] and its association with increased risk of injury [22], there is a need to address the isometric strength testing of hand and forearm muscles of sit-skiers before and immediately after a training session.

Under these circumstances, this case study aims to inform about the physiological laboratory evaluation and the TL experienced by a sit-skier during adaptive slalom waterskiing training sessions aimed at improving the DWS technique. A secondary objective is to assess the effects of these training sessions on handgrip strength.

## Materials and methods

### Participant

The athlete who took part in this case study was a 28-year-old male (height: 1.77 m, body mass: 55.0 kg) with paraplegia (complete injury at T5, ASIA Impairment Scale A) [23] as a result of a motor vehicle accident that occurred 7 years ago. He had 3 years of adaptive waterskiing experience. He mostly practiced this sport at a cable park and during open waterskiing on a reservoir, adopting a sitting position with the knees higher than the hips at a tow rope length of 18.2 m. He was considered an intermediate-advanced skier [5,7], and typically struggled to get out of the water without assistance, resulting in a failed or thoroughly exhausting attempt. He decided to undergo training sessions to be able to get on top of the water independently by learning a new DWS technique. He received a detailed explanation of the study and signed the informed consent procedure. The study was approved by the Ethics Committee of the University of León, Spain and conducted in accordance with the Declaration of Helsinki.

### Pretraining laboratory test

#### Cardiorespiratory fitness

The participant’s cardiorespiratory fitness was identified through a series of laboratory tests completed two days before the training program started. An incremental test was performed on an arm-crank ergometer with frictional load (Monark Rehab Trainer 881 E, Monark Exercise AB, Varberg, Sweden). The test started at 5 W and the power output was manually increased by 5 W every 1 min until the participant could not maintain a pedaling frequency ≥ 50 rev·min^−1^. The HR response was measured telemetrically every 5 s (Polar Team System 2, Polar Electro Oy, Kempele, Finland), ECG and respiratory gas exchange were continuously measured (Medisoft Ergocard, Medisoft Group, Sorinnes, Belgium). End criteria for reaching maximal oxygen uptake (VO_2max_) were achieved: VO_2_ plateau (≤ 150 mL·min^−1^ with increasing workload), maximal HR equivalent to ± 10 beats·min^−1^ of age-predicted maximal HR for upper body exercise (200 - age) [24] and a RER ≥ 1.10 [25]. The ventilatory thresholds were identified separately by two researchers [26], and both researchers agreed.

#### Pulmonary function

Pulmonary function tests were conducted following standard protocols published by the American Thoracic Society/European Respiratory Society [27]. The participant performed the tests in his own wheelchair and wearing a nose clip. The forced vital capacity (FVC), the forced expiratory volume in one second (FEV1), and slow vital capacity (SVC) were measured using the MIR Spirobank I® portable electronic spirometer coupled to a laptop computer with spirometry software WinspiroPRO® 1.3 (Medical International Research, Rome, Italy).

### Training sessions

The training program consisted of six sessions (3 daily sessions per week during two consecutive weeks) aimed at learning, practicing, and improving two DWS techniques. The traditional technique implies that the skier is sitting in the water and lets the towboat pull him up on top of the water without assistance [5]. The alternative assisted technique uses “the boom” (training bar) placed on the port side of the towboat^9^, and a short rope attached to it that is eventually extended so that the sit-skier becomes aligned in the centre of the wakes and behind the towboat.

The session started with a standardized 5 min warm up consisting of mobility exercises in the towboat. After that, the alternative technique was introduced to begin the training session on water. Once a successful DWS was achieved, the sit-skier practiced typical activities, mainly a series of slalom runs splitting his time between open water (out of the slalom course) and the inner slalom, an adapted course with buoys at 6.4 m from the axis of the slalom course [28]. If the sit-skier fell during a run, he had to complete a DWS and restart the practice with a rest period between, allowing coaching feedback. The training program was carried out at a reservoir in the north of Spain.

Special considerations were applied for the participant since spinal cord lesion at T5 situated him at increased risk for hypothermia and autonomic dysreflexia: a) he was asked to empty the bladder or the urine collection bag before each practice; b) the use of a wetsuit was used as preventative measure during practices; c) hydration ad libitum was provided; d) and prolonged rest periods in the towboat with wet clothes were avoided [29]. Weather conditions were variable across training sessions, with air temperatures between 20–32°C, wind speeds between 12.0–37.0 km·h^−1^, and reservoir water temperatures between 14–16°C. Skiing was performed in sheltered parts of the reservoir to reduce wind exposure and use flatter water, away from another boat's wakes, with the sit-skier wearing a waterskiing vest [30]. To help minimize the variables that might influence the outcomes, all of the training sessions were done at similar times throughout the study period (from 11:45 a.m. to 2:15 p.m.), using the same towboat and with the same experienced towboat driver setting the towboat speed with PerfectPass speed control (PerfectPass Control Systems Inc., Dartmouth, NS, Canada), on the same inner-slalom course, and without change in length of the tow rope.

#### Internal training load

The exercise demands were quantified on the basis of HR and RPE [15,16]. HR was recorded every 5 s during each training session (Polar Team System 2, Polar Electro Oy, Kempele, Finland). After the training sessions, HR data was downloaded to a computer through a specific software (Polar Pro Trainer 5, Polar Electro Oy, Kempele, Finland). The HR response was categorized into 3 intensity zones according to the HR values corresponding to the ventilatory (VT) and respiratory compensation (RCT) thresholds [16]: zone 1 (low-intensity exercise), below VT; zone 2 (moderate-intensity exercise), between VT and RCT; and zone 3 (high-intensity exercise), above RCT. These zones were used to calculate the TL by multiplying the time spent in zone 1, 2 and 3 by the constants 1, 2, and 3, respectively, and the TL score being obtained by summating the results of the 3 phases [16]. The participant’s session rating of perceived exertion (sRPE) was obtained using the modified Borg (0–10) scale collected ~30 min after each training session [15]. The participant was already familiar with this scale since it was routinely used to control the training intensity during the year before starting the study. Training sessions were divided into 3 intensity levels according to the sRPE: moderate (sRPE < 5), hard (sRPE 5–6), and fairly hard (sRPE > 6) [31]. Training load was calculated by multiplying the sRPE value by the duration of the training session [15].

#### Handgrip strength

Handgrip strength was measured with a digital dynamometer (Takei TKK 5401 Grip-D, Tokyo, Japan), immediately before and after the training sessions. A standardized testing position was adopted following the procedure of the American Society of Hand Therapists (ASHT), with the participant in a sitting position in the towboat and the elbow bent at 90° [32]. The participant was able to self-select the handgrip position on the dynamometer with which he could exert the highest force [33]. Peak force was measured in kg during a 5 s period; a force of 3 consecutive maximal repetitions was recorded for the dominant hand. There were 60 s rest intervals between repetitions. The greatest value produced during the three repetitions was used for analysis. The strength decrement index was calculated on each training session: 100% × (Initial Max–Final Max) × Initial Max [34].

All measurements were conducted by the same researcher. He had sufficient experience as a coach of athletes with physical disabilities and was familiar with the procedures conducted to evaluate physical conditions in this population. To quantify characteristics and events during training sessions, all activities were observed by this expert who made written recordings about their nature and duration, as well as towboat speeds (Traceable manual digital chronometer VWR, Pennsylvania, USA). Activities were categorized into four broad groups depending on their nature: *deep-water start*, included proper position of the body while sitting in the water to start from a resting position until the skier planed on top of the water and began a run; *open-water practice*, recorded from the time the skier started to being pulled behind the towboat, immediately after the DWS, to the moment the skier fell, released the tow rope handle, and sank into the water, or effectively entered the inner-slalom course; *inner-slalom course practice*, recorded from the time the skier began waterskiing through the course to right before he either fell or went back to open waterskiing; and *rest period*, considered all situations in which the skier was stationary on the water receiving coaching feedback or recovering and getting back on the sit-ski after falling. Within the *deep-water start* category, further differentiation was made to identify between *alternative* (ADWS) and *traditional* (TDWS). The number of completed DWSs where the skier was successful, as well as the variation range of skier’s speed, were registered.

### Statistical analyses

Data were expressed as mean ± standard deviation (SD). The assumption of normality was verified using the Shapiro-Wilk test. The relationship between HR-based TL method and RPE-based TL method was determined by means of Pearson’s correlation coefficient (*r*). The magnitude of the correlation (*r*) was defined as: trivial (< 0.1), small (0.1–0.3), moderate (0.3–0.5), large (0.5–0.7), very large (0.7–0.9) and nearly perfect (> 0.9) [35]. Values of p < .05 were considered statistically significant. A paired student's t‐test was used to compare pre- and post-training handgrip strength values. The statistical analyses were performed using the Statistical Package for the Social Sciences (SPSS version 24.0, Chicago, Illinois, USA).

## Results

### Pretraining laboratory test

#### Cardiorespiratory fitness

The VO_2max_, maximal power output and HR were 22 ml·kg^−1^·min^−1^, 65 W and 182 beats·min^−1^, respectively. The VT was identified at 47% VO_2max_ at a power output and HR of 25 W and 143 beats·min^−1^, respectively. The RCT was found at 72% VO_2max_ at a power output and HR of 40 W and 163 beats·min^−1^, respectively.

#### Pulmonary function

The pulmonary function parameters measured were: FVC, 3.49 L; FEV1, 3.40 L; and VC, 3.23 L.

### Training sessions

The main characteristics of the training sessions are shown in Table 1. Average session duration was 30 ± 7 min. The ADWS phase made up 8.4 ± 5.2% of the total time spent during a session, TDWS 5.5 ± 4.4%, *open-water practice* 48.3 ± 23.6%, *inner-slalom course practice* 16.3 ± 16.5%, and *rest period* 21.5 ± 13.7. Towboat speed ranged between 31–43 km·h^−1^ throughout the training sessions. During the 6-day training period, a total of 23 DWSs were undertaken. A greater length of time was spent with the alternative method (total attempts = 13, total time = 16 min 6 s, mean time per attempt = 74 s, completion rate = 100%) compared to the traditional method (total attempts = 10, total time = 10 min 13 s, mean time per attempt = 61 s, completion rate = 20%).

**Table 1.** Summary of data for the training sessions.

| Training session | 1 | 2 | 3 | 4 | 5 | 6 |
| --- | --- | --- | --- | --- | --- | --- |
| Total duration (min:s) | 34:33 | 27:43 | 29:50 | 40:46 | 28:31 | 20:00 |
| Alternative deep-water start | 03:19 | 01:26 | 05:10 | 04:16 | 01:05 | 00:50 |
| Traditional deep-water start | 03:25 | - | 02:46 | 00:55 | 02:37 | 00:30 |
| Open-water practice | 17:05 | 26:17 | 11:31 | 14:29 | 08:47 | 08:05 |
| Inner-slalom course practice | 03:44 | - | - | 06:20 | 08:58 | 08:00 |
| Rest period | 07:00 | - | 10:23 | 14:46 | 07:04 | 02:35 |
| **Towboat speed (km·h<sup>-1</sup>)** | 31 | 34 | 34 | 34 | 34-43 | 37-40 |
| **Alternative deep-water start** |  |  |  |  |  |  |
| No. of attempts | 2 | 1 | 5 | 3 | 1 | 1 |
| No. of successful attempts | 2 | 1 | 5 | 3 | 1 | 1 |
| **Traditional deep-water start** |  |  |  |  |  |  |
| No. of attempts | 2 | - | 2 | 2 | 3 | 1 |
| No. of successful attempts | 0 | - | 0 | 2 | 0 | 0 |

#### Internal training load

Physiological demands experienced by the sit skier are shown in Table 2. When HR data were analyzed using percentages of maximal HR (%HR_max_), ADWS phases were performed at an average HR of 70.8 ± 6.3% HR_max_. During TDWS phases, HR reached an average of 69.6 ± 4.5% HR_max_. The average intensity during all sessions was 71.8 ± 4.4% HR_max_. The participant spent 76.7 ± 15.3, 21.7 ± 14.2, and 1.7 ± 1.6% of total time in zone 1, 2 and 3, respectively. Mean sRPE across training sessions was 4.7 ± 1.6. The mean TL assessed according to the HR and sRPE was 39.6 ± 3.1 arbitrary units (AU) and 151.6 ± 61.1 AU, respectively. A non-significant moderate positive correlation of *r* = .45 (p = .37) was observed between HR-based TL and sRPE-based TL.

**Table 2.**
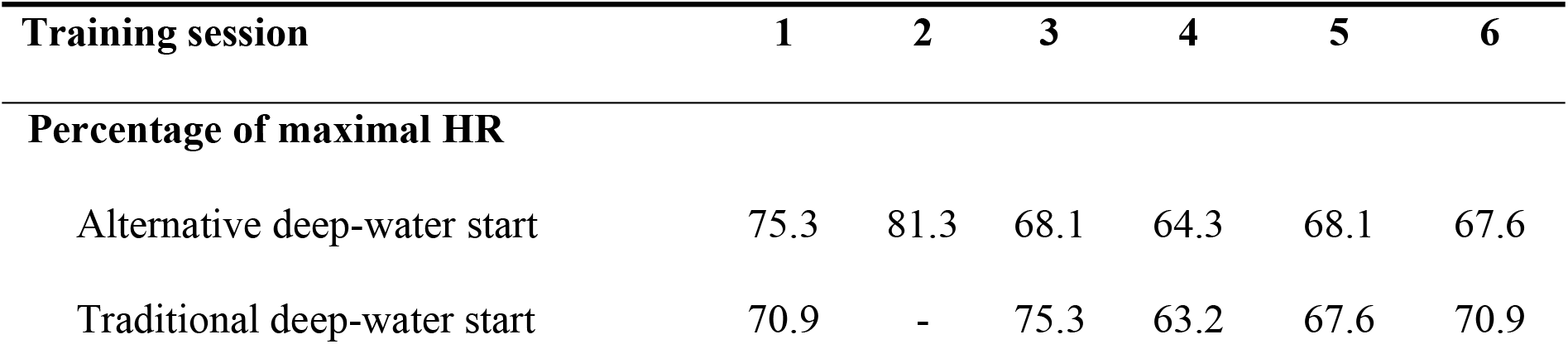
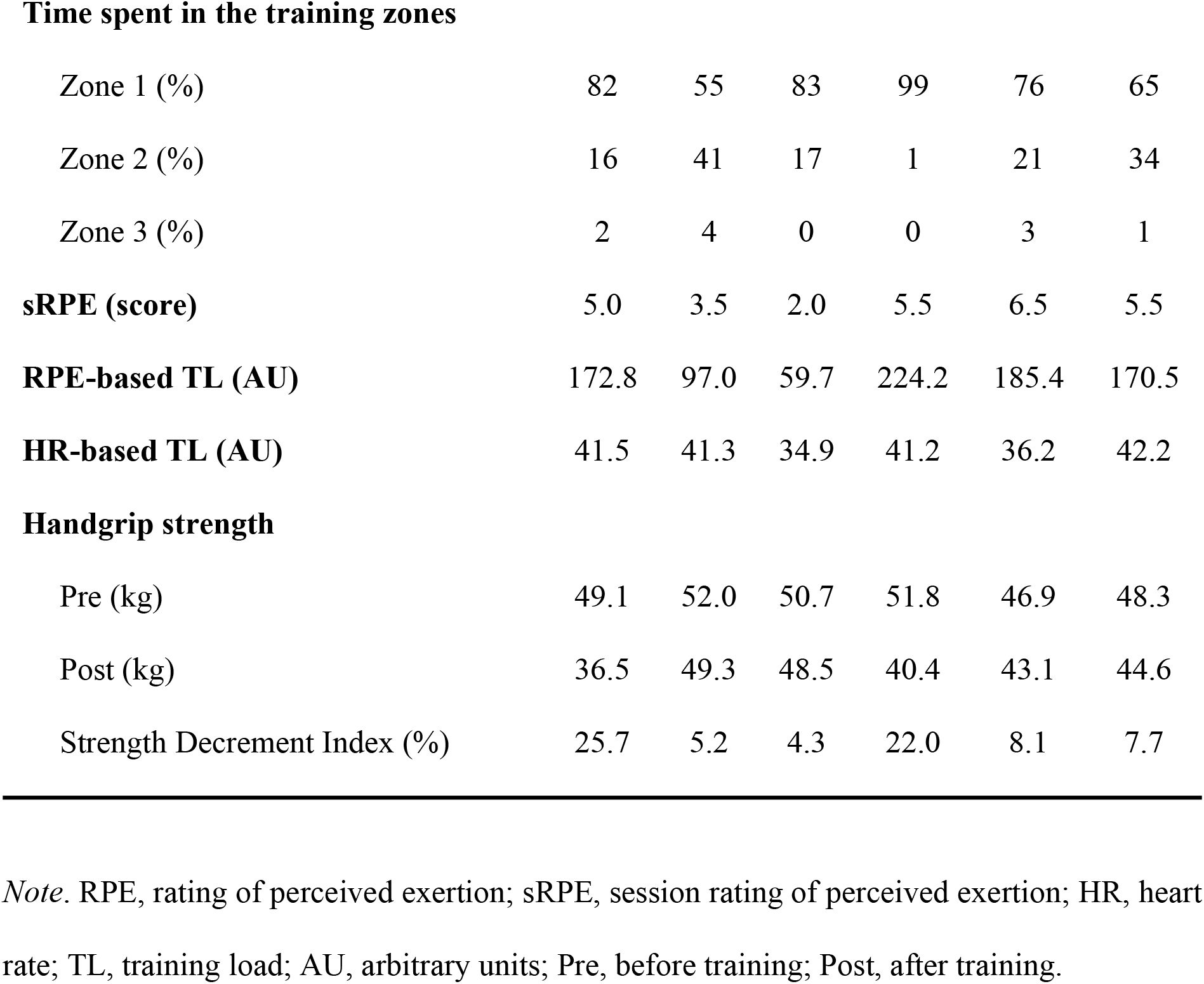
Percentage of maximal heart rate during deep-water starts and percentage time spent in each training zone, session rating of perceived exertion, training load, pre- and post-training values on handgrip strength measurements and Strength Decrement Index during the training sessions.

| Training session | 1 | 2 | 3 | 4 | 5 | 6 |
| --- | --- | --- | --- | --- | --- | --- |
| **Percentage of maximal HR** |  |  |  |  |  |  |
| Alternative deep-water start | 75.3 | 81.3 | 68.1 | 64.3 | 68.1 | 67.6 |
| Traditional deep-water start | 70.9 | - | 75.3 | 63.2 | 67.6 | 70.9 |
| **Time spent in the training zones** |  |  |  |  |  |  |
| Zone 1 (%) | 82 | 55 | 83 | 99 | 76 | 65 |
| Zone 2 (%) | 16 | 41 | 17 | 1 | 21 | 34 |
| Zone 3 (%) | 2 | 4 | 0 | 0 | 3 | 1 |
| sRPE (score) | 5.0 | 3.5 | 2.0 | 5.5 | 6.5 | 5.5 |
| RPE-based TL (AU) | 172.8 | 97.0 | 59.7 | 224.2 | 185.4 | 170.5 |
| HR-based TL (AU) | 41.5 | 41.3 | 34.9 | 41.2 | 36.2 | 42.2 |
| **Handgrip strength** |  |  |  |  |  |  |
| Pre (kg) | 49.1 | 52.0 | 50.7 | 51.8 | 46.9 | 48.3 |
| Post (kg) | 36.5 | 49.3 | 48.5 | 40.4 | 43.1 | 44.6 |
| Strength Decrement Index (%) | 25.7 | 5.2 | 4.3 | 22.0 | 8.1 | 7.7 |

*Note.* RPE, rating of perceived exertion; sRPE, session rating of perceived exertion; HR, heart rate; TL, training load; AU, arbitrary units; Pre, before training; Post, after training.

#### Handgrip strength

The pre- and post-training handgrip strength values of the participant are also shown in Table 2. The mean post-training value (43.7 ± 4.9 kg) was significantly lower (p = .024) than pre-training value (49.8 ± 2.0 kg). Strength decrement index mean values, calculated as the deterioration of strength as a proportion of pretraining values, were 12.2 ± 9.2 %. Strength decrement index values were over 20% for the first and the fourth session.

## Discussion

This study aimed to provide data about the physiological laboratory evaluation and the TL experienced by a sit-skier with paraplegia while practicing slalom training sessions, with a direct measurement of the cardiovascular, perceptual, and handgrip strength imposed on skiers during DWS and subsequent phases of slalom waterskiing.

There is little published data on the physiological evaluation of adaptive waterskiers, or indeed able-bodied waterskiers. Peak oxygen uptake for our participant was similar to that observed for a female waterskier of the sitting class (25 ml◻kg^−1^◻min^−1^) [36]. The lower values compared to those reported for a group of male standing elite waterskiers (54.5 ± 6.2 ml·kg^−1^·min^−1^) have been very likely caused by reduced active muscle mass during exercise in laboratory [37]. Additionally, the pulmonary function values observed for the sit-skier were slightly below those values reported in sixteen highly trained wheelchair basketball players experiencing mostly SCI (13 out of 16) [38]. We believe this occurs because our sit-skier may experience loss of pulmonary function due to the high level of paraplegia, with 81% of predicted FVC at the T5 level [39].

Very few studies have investigated the use of HR in athletes with paraplegia as a means of monitoring exercise intensity [3]. According to the data registered, our participant spent most of his training practice in zone 1 or zone 2 in all recorded sessions. This finding suggests the predominance of aerobic metabolism when practicing adaptive waterskiing in a sitting position, in accordance with standing slalom waterskiing being described as dependent on aerobic metabolism [14]. This has also been the case for other sport modalities such as Paralympic alpine sit-ski [40], where it appears that sit-skiing athletes experience a moderate strain even under repeated race-like situations. In this regard, it has been suggested that compared to standing alpine skiers, endurance capacities play a smaller role for sit-skiers [40]. It is to be expected, therefore, that the same applies to standing and sitting slalom waterskiing, since moderate-to-high levels of both anaerobic and aerobic power among elite standing waterskiers have been found [11,12].

In addition, our research observed for the first time HR values expressed as %HR_max_ during DWS, ranging from 63.2% to 81.3%. These results are likely to be related to the high static (related to blood pressure) and low dynamic demands (in terms of cardiac output) of waterskiing [41]. The participant’s %HR_max_ values during DWS are comparable to previous reports of Paralympic alpine skiing sitting athletes (from 67.0% to 83.0%) [40]. A possible explanation for this might be that the duration of the giant slalom competition-like runs were approximately 55 s, showing a nature of short duration similar to that found for the DWS in our participant. Similar HR values have also been observed in sports such as wheelchair tennis or wheelchair basketball during practice or match play [42–45]. These similarities may partly be explained by the fact that both adaptive slalom waterskiing and the aforementioned wheelchair sports present an intermittent nature.

To the best of our knowledge, this is the first attempt to investigate the sRPE in both standing and sitting slalom waterskiing. For TL monitoring in the population with SCI, sRPE is one of the most practical methods [3]. The adaptive slalom waterskiing training sessions in this study ranged from moderate to fairly hard intensity according to sRPE values. It must be noted that the perceptual response to training sessions offered values greater than 5 on four occasions, suggesting that our participant was experiencing slalom sit-skiing as a hard activity in general. The yields in this investigation were slightly higher compared to those of four highly trained international WT players during singles matches, where the mean of the sRPE value was described as *somewhat* hard [46]. This discrepancy could be attributed to differences in the levels of ability and experience between our sit-skier and these WT players. Previous research has demonstrated that individuals’ RPE may be influenced by skiing skills, where recreational alpine skiers with advanced skiing ability report lower RPE values [47].

It is now well established that monitoring of TL can provide both valuable information about an athlete’s adaptation to a training program and support for coaches in the design of appropriate training programs [48]. The current study found that HR and sRPE-based methods of TL showed different scores. The obtained TL values were lower than those observed in highly trained WB players (55.3–67.5 AU) during a small-sided games session [49]. Also, greater values of TL using the same HR-based method that we used in this study have been provided during court-based training in elite wheelchair rugby (WR) athletes (247 ± 74 AU) [50]. A note of caution is due here since these greater values could be due to the differences between sport modalities (WB and WR present typically high intensity intermittent activities involving a wheelchair) and the different injury characteristics in participants (WB includes heterogeneous disabilities, and WR is performed by athletes with cervical SCI). Regarding the sRPE-based method for quantifying TL, our values are far above those observed in wheelchair basketball for a small-sided games session (99.3 ± 26.9 AU) [49]. On the other hand, higher sRPE-based TL values (934 ± 359 AU) during WR practice were documented [50]. Again, these comparisons must be interpreted with caution because the aforementioned studies were different in terms of participants, activities, and duration of training sessions.

The limited availability of athletes with a SCI participating in research studies explain why various training recommendations are given on an individual basis or according to coaches’, athletes’ and scientist’s personal experiences [51]. In this context, little is known with respect to the relationship between objective and subjective TL on athletes with disabilities. In our study, we identified a moderate but not significant correlation between the HR and sRPE-based TL (*r* = .45; p = .37). These results are likely to be related to the large static component of waterskiing activities. It has been shown that during fatiguing isometric exercise, for paraplegic patients there is only a modest increase in HR since this index appears to be centrally mediated [52]. With regard to RPE, it might be that both the repetition of DWS and the longer duration of training sessions when compared to regular individual ski time of ~15 min in slalom courses [14] could lead to a greater accumulation of muscle fatigue. The small alterations in force at the precise moment along with the extremely large forces to react to sudden external disturbances when performing highly skilled and well‐rehearsed moves, typical in slalom waterskiing [14], could also contribute to peripheral fatigue [53]. Therefore, the sRPE score as an indicator of exercise intensity may be affected due to its psychobiological nature that integrates various information including the signals produced by the peripheral working muscles [54]. The findings of the current study are in line with previous research reporting only small or trivial non-significant correlations between HR and sRPE-based TL in relation to WB small-sided games (range r = − .30; ± .27 to .26; ± .28; p >.05) [49]. Moderate but significant relationships between HR-based and RPE-based methods have been also found to quantify match load in WB (r = .63 – .65, p < .001) [55]. In contrast, a very large correlation was found between variables (r = .81) in WR training sessions [50]. It may be the case, therefore, that these variations are caused by the differences in the studies concerning levels of functional ability among participants or the variability in the conditions recorded (small-sided games, conventional training sessions and matches).

In the present study, we found a statistically significant reduction in the handgrip strength of the participant once the sessions were finished. This could be due to the fact that intrinsic and extrinsic finger flexors, as well as forearm extensors, are very active while gripping the tow rope handle during most of the practice time [14]. In this regard, in other adaptive sports modalities where this musculature is heavily involved, such as judo, similar reductions on handgrip strength were observed [56]. These results support previous research indicating that waterskiing is a high-demand sport in terms of grip strength [22]. It should be noted that waterskiing technique can have an impact on forearm muscle loading during gripping [22]. Bearing all of the above in mind, to maintain optimum levels of neuromuscular performance during adaptive slalom waterskiing, we recommend that coaches focus on optimizing the position of upper limbs during practice to minimize handgrip force.

In spite of its originality, there are two important limitations that should be taken into account when interpreting the obtained results. In the first place, the data comes from one participant. Thus, extrapolation to other adaptive athletes is limited. Secondly, the registered data is somehow incomplete since we did not perform a time-motion analysis addressing the waterskier’s external load (i.e. distance, speed, duration). This fact prevents carrying out a more thorough analysis regarding the TL experienced by the participant.

## Conclusions

In conclusion, performing adaptive slalom waterskiing training by a sit-skier with paraplegia involved in the present study a moderate physiological internal load, which was perceived as a hard effort. Moreover, the significant decrement in handgrip strength offers valuable data about the extent of the grip-strength contribution required in a sitting position. These findings provide novel information regarding the TL that can be experienced while practicing this sport modality.

## Acknowledgments

The authors would like to acknowledge the considerable contributions while collecting data of CD Esquí Náutico León coaches and skiers, and also the institutional support from the CRE Discapacidad y Dependencia (San Andrés del Rabanedo), Institute for Older Persons and Social Services (IMSERSO). This work was supported by a grant from the Ministry of Education, Culture and Sports, Government of Spain awarded to David Suárez-Iglesias [FPU12/05828].

## References

1. Leprêtre P-M, Goosey-Tolfrey VL, Janssen TWJ, Perret C. Editorial: Rio, Tokyo Paralympic Games and Beyond: How to Prepare Athletes with Motor Disabilities for Peaking. Front Physiol. 2016;7:497.

2. Baumgart JK, Brurok B, Sandbakk Ø. Peak oxygen uptake in Paralympic sitting sports: A systematic literature review, meta- and pooled-data analysis. Bragazzi N, editor. PLoS One. 2018;13(2):e0192903.

3. Simim MAM, de Mello MT, Silva BVC, Rodrigues DF, Rosa JPP, Couto BP, et al. Load monitoring variables in training and competition situations: A systematic review applied to wheelchair sports. Adapt Phys Act Q. 2017;34(4):466–83.

4. Mulhollon S, Casey J. Adaptive sports and equipment for veterans with spinal cord injuries: community partnerships are key to a year-round adapted sports programs for Milwaukee veterans. Palaestra. 2016;30(4):20–4.

5. Bray-Miners J, Runciman RJ, Monteith G. Water skiing biomechanics: a study of advanced skiers. Proc Inst Mech Eng Part P J Sport Eng Technol. 2012;227(2):137–46.

6. Keverline JP, Englund R, Cooney TE. Takeoff forces transmitted to the upper extremity during water-skiing. Orthopedics. 2003 Jul;26(7):707–10.

7. Runciman RJ. Water-skiing biomechanics: a study of intermediate skiers. Proc Inst Mech Eng Part P J Sport Eng Technol. 2011;225(4):231–9.

8. USA Water Ski. Level 1 Instructor’s Manual. A Self-Study Course For Learning How To Teach Beginning Water Skiing. p. 72. Available from: http://www.bebercamp.com/images/uploads/pdfs/5_-_level_1-_manual.pdf

9. Kegel B. Physical fitness. Sports and recreation for those with lower limb amputation or impairment. J Rehabil Res Dev Clin Suppl. 1985;(1):1–125.

10. Bray-Miners J, Runciman RJ, Monteith G, Groendyk N. Biomechanics of slalom water skiing. Proc Inst Mech Eng Part P J Sport Eng Technol. 2015;229(1):47–57.

11. Leggett SH, Fulton MN, Pollock ML, Carpenter DM, Graves JE, Shank MB, et al. Physiological evaluation of professional water-skiers. J Strength Cond Res. 1994;8(1):20–7.

12. Leggett SH, Kenney K, Eberhardt T. Applied Physiology of Water-Skiing. Sports Med. 1996;21(4):262–76.

13. Eberhardt TPT. Sports performance series: an anatomical description of the slalom turn in water skiing and how to condition for this sport. Strength Cond J. 1988;10(6):4–9.

14. Mullins NM. Slalom water skiing: physiological considerations and specific conditioning. Strength Cond J. 2007;29(4):42–54.

15. Foster C, Florhaug JA, Franklin J, Gottschall L, Hrovatin LA, Parker S, et al. A new approach to monitoring exercise training. J Strength Cond Res. 2001;15(1):109–15.

16. Rodríguez-Marroyo JA, Villa G, García-López J, Foster C. Comparison of heart rate and session rating of perceived exertion methods of defining exercise load in cyclists. J Strength Cond Res. 2012;26(8):2249–57.

17. Yanci J, Granados C, Otero M, Badiola A, Olasagasti J, Bidaurrazaga-Letona I, et al. Sprint, agility, strength and endurance capacity in wheelchair basketball players. Biol Sport. 2015;32(1):71–8.

18. Porto YC, Almeida M, de Sá CC, Schwingel PA, Zoppi CC. Anthropometric and physical characteristics of motor disabilited paralympic rowers. Res Sport Med. 2008;16(3):203–12.

19. Iturricastillo A, Granados C, Yanci J. Changes in body composition and physical performance in wheelchair basketball players during a competitive season. J Hum Kinet. 2015;48:157–65.

20. Oliveira L, Oliveira S, Guimarães F, Costa M. Contributions of body fat, fat free mass and arm muscle area in athletic performance of wheelchair basketball players. Motricidade. 2017;13(2):36.

21. Rudolph L, Willick S, Teramoto M, Cushman DM. Adaptive Sports Injury Epidemiology. Sports Med Arthrosc. 2019;27(2):e8–11.

22. Rosa D, Di Donato SL, Balato G, D’Addona A, Schonauer F. Supinated forearm is correlated with the onset of medial epicondylitis in professional slalom waterskiers. Muscles Ligaments Tendons J. 2016;6(1):140–6.

23. Kirshblum SC, Burns SP, Biering-Sorensen F, Donovan W, Graves DE, Jha A, et al. International standards for neurological classification of spinal cord injury (revised 2011). J Spinal Cord Med. 2011;34(6):535–46.

24. Smith PM, Price MJ. Upper-body exercise. In: Edward M. Winter, Andrew M. Jones, R.C. Richard Davison, Paul D. Bromley THM, editor. Sport and Exercise Physiology Testing Guidelines: Volume I - Sport Testing - The British Association of Sport and Exercise Sciences Guide [Internet]. 1st ed. London : Routledge; p. 158–64.

25. Midgley AW, McNaughton LR, Polman R, Marchant D. Criteria for determination of maximal oxygen uptake: a brief critique and recommendations for future research. Sports Med. 2007;37(12):1019–28.

26. Au JS, Sithamparapillai A, Currie KD, Krassioukov AV., MacDonald MJ, Hicks AL. Assessing ventilatory threshold in individuals with motor-complete spinal cord injury. Arch Phys Med Rehabil. 2018;99(10):1991–7.

27. Miller MR, Hankinson J, Brusasco V, Burgos F, Casaburi R, Coates A, et al. Standardisation of spirometry. Eur Respir J. 2005;26(2):319–38.

28. International Waterski & Wakeboard Federation. 2019 Technical rules for water ski for the disabled. 2019. Available from: http://iwwfed.com/2017-technical-rules-water-ski-for-the-disabled/

29. De Luigi AJ. Adaptive Sports Medicine. 1st ed. De Luigi AJ, editor. Champaign, IL: Springer International Publishing; 2018. 402 p.

30. Loughlin S. Investigation of injuries occurring within competitive waterskiing in the UK. Int J Exerc Sci. 2013;6(1):29–42.

31. Rodríguez-Marroyo JA, Antoñan C. Validity of the session rating of perceived exertion for monitoring exercise demands in youth soccer players. Int J Sports Physiol Perform. 2015;10(3):404–7.

32. Sisto SA, Dyson-Hudson T. Dynamometry testing in spinal cord injury. J Rehabil Res Dev. 2007;44(1):123–36.

33. Boadella JM, Kuijer PP, Sluiter JK, Frings-Dresen MH. Effect of self-selected handgrip position on maximal handgrip strength. Arch Phys Med Rehabil. 2005;86(2):328–31.

34. Reuter SE, Massy-Westropp N, Evans AM. Reliability and validity of indices of hand-grip strength and endurance. Aust Occup Ther J. 2011;58(2):82–7.

35. Hopkins WG. Probabilities of clinical or practical significance. Sportscience. 2002;(6). Available from: http://www.sportsci.org/jour/0201/wghprob.htm

36. Goosey-Tolfrey V, Tolfrey K. The oxygen uptake-heart rate relationship in trained female wheelchair athletes. J Rehabil Res Dev. 2004;41(3B):415–20.

37. Goosey-Tolfrey V, Leicht CA. Field-based physiological testing of wheelchair athletes. Sport Med. 2013;43(2):77–91.

38. Goosey-Tolfrey V, Foden E, Perret C, Degens H. Effects of inspiratory muscle training on respiratory function and repetitive sprint performance in wheelchair basketball players. Br J Sports Med. 2010;44(9):665–8.

39. Linn W, Spungen A, Gong, Jr H, Adkins R, Bauman A, Waters R. Forced vital capacity in two large outpatient populations with chronic spinal cord injury. Spinal Cord. 2001;39(5):263–8.

40. Goll M, Wiedemann MSF, Spitzenpfeil P. Metabolic demand of paralympic alpine skiing in sit-skiing athletes. J Sports Sci Med. 2015;14(4):819–24.

41. Mitchell JH, Haskell W, Snell P, Van Camp SP. Task Force 8: classification of sports. J Am Coll Cardiol. 2005;45(8):1364–7.

42. Barfield JP, Malone LA, Coleman TA. Comparison of heart rate response to tennis activity between persons with and without spinal cord injuries: implications for a training threshold. Res Q Exerc Sport. 2009;80(1):71–7.

43. Croft L, Dybrus S, Lenton J, Goosey-Tolfrey V. A comparison of the physiological demands of wheelchair basketball and wheelchair tennis. Int J Sports Physiol Perform. 2010;5(3):301–15.

44. Roy JLP, Menear KS, Schmid MMA, Hunter GR, Malone LA. Physiological responses of skilled players during a competitive wheelchair tennis match. J Strength Cond Res. 2006;20(3):665–71.

45. Sindall P, Lenton JP, Tolfrey K, Cooper RA, Oyster M, Goosey-Tolfrey VL. Wheelchair tennis match-play demands: effect of player rank and result. Int J Sports Physiol Perform. 2013;8(1):28–37.

46. Sánchez-Pay A, Torres-Luque G, Sanz-Rivas D. Match activity and physiological load in wheelchair tennis players: a pilot study. Spinal Cord. 2016;54(3):229–33.

47. Scheiber P, Krautgasser S, von Duvillard SP, Müller E. Physiologic responses of older recreational alpine skiers to different skiing modes. Eur J Appl Physiol. 2009;105(4):551–8.

48. Halson SL. Monitoring training load to understand fatigue in athletes. Sports Med. 2014;44 Suppl 2(Suppl 2):S139–47.

49. Iturricastillo A, Granados C, Los Arcos A, Yanci J. Objective and subjective methods for quantifying training load in wheelchair basketball small-sided games. J Sports Sci. 2017;35(8):749–55.

50. Paulson TAW, Mason B, Rhodes J, Goosey-Tolfrey VL. Individualized internal and external training load relationships in elite wheelchair rugby players. Front Physiol. 2015;6:388.

51. Perret C. Elite-adapted wheelchair sports performance: a systematic review. Disabil Rehabil. 2017;39(2):164–72.

52. Petrofsky J. Blood pressure and heart rate response to isometric exercise: the effect of spinal cord injury in humans. Eur J Appl Physiol. 2001;85(6):521–6.

53. Ferguson RA. Limitations to performance during alpine skiing. Exp Physiol. 2010;95(3):404–10.

54. Borg GA. Psychophysical bases of perceived exertion. Med Sci Sports Exerc. 1982;14(5):377–81.

55. Iturricastillo A, Yanci J, Granados C, Goosey-Tolfrey V. Quantifying wheelchair basketball match load: a comparison of heart-rate and perceived-exertion methods. Int J Sports Physiol Perform. 2016;11(4):508–14.

56. Bonitch-Góngora JG, Bonitch-Domínguez JG, Padial P, Feriche B. The effect of lactate concentration on the handgrip strength during judo bouts. J Strength Cond Res. 2012;26(7):1863–71.

